# Knowledge Beacons: Web Service Workflow for FAIR Data Harvesting of Distributed Biomedical Knowledge

**DOI:** 10.1101/2020.04.06.027425

**Authors:** Lance M. Hannestad, Vlado Dančík, Meera Godden, Imelda W. Suen, Kenneth C. Huellas-Bruskiewicz, Benjamin M. Good, Christopher J. Mungall, Richard M. Bruskiewich

## Abstract

The continually expanding distributed global compendium of biomedical knowledge is diffuse, heterogeneous and huge, posing a serious challenge for biomedical researchers in knowledge harvesting: accessing, compiling, integrating and interpreting data, information and knowledge. In order to accelerate research towards effective medical treatments and optimizing health, it is critical that efficient and automated tools for identifying key research concepts and their experimentally discovered interrelationships are developed.

As an activity within the feasibility phase of a project called “Translator” **(https://ncats.nih.gov/translator**) funded by the National Center for Advancing Translational Sciences (NCATS) to develop a biomedical science knowledge management platform, we designed a Representational State Transfer (REST) web services Application Programming Interface (API) specification, which we call a Knowledge Beacon. Knowledge Beacons provide a standardized basic workflow for the discovery of concepts, their relationships and associated supporting evidence from distributed online repositories of biomedical knowledge. This specification also enforces the annotation of knowledge concepts and statements to the NCATS endorsed the Biolink Model data model and semantic encoding standards (**https://biolink.github.io/biolink-model/**). Implementation of this API on top of diverse knowledge sources potentially enables their uniform integration behind client software which will facilitate research access and integration of biomedical knowledge.

**Availability:** The API and associated software is open source and currently available for access at **https://github.com/NCATS-Tangerine/translator-knowledge-beacon**.

## Introduction

A serious challenge to impactful biomedical research is the one that biomedical researchers encounter when identifying and accessing pertinent information: the diffuse and voluminous nature of such data and knowledge. The large, rapidly growing compendium of published scientific literature is characterized by diverse data encoding standards; numerous, distinct, heterogeneous, large and often siloed public research data repositories; relatively inaccessible health records; numerous clinical trial and adverse event reports, all spread across disease communities and biomedical disciplines. The current distributed nature of this knowledge and associated (meta-)data silos impedes the discovery of related concepts and the relationships between them, an activity one might call “Knowledge Harvesting”. Many efforts to overcome this challenge focus on data management principles to make such resources “**F**indable, **I**nteroperable, **A**ccessible and **R**eusable” (FAIR) [1,2].

Web access to bioinformatics data spans many generations of web service standards tagged with many acronyms, e.g. CORBA [3], SOAP/BioMOBY [4] and SADI [5], the latter an exemplar of the more general paradigm of “Linked Open Data” using OWL/RDF and SPARQL technology, including Linked Open Fragments [6].

A popular web service standard currently in use is the Swagger 2.0 or OpenAPI 3.0 specified REST API (https://github.com/OAI). Many extant online biomedical data sources currently provide such REST API implementations for accessing their data. API registries exist to index such APIs to facilitate access (for example, the Smart API Registry; https://smart-api.info/) and generalized tools are available to explore the space of such web services (notably, the Biothings API and Explorer; https://biothings.io/). However, the heterogeneity of such APIs can be a barrier to efficient biomedical knowledge integration.

Here we present a REST-based web services specification called the Knowledge Beacon API (Beacon API) that enables a basic workflow for the discovery of, and navigation through, biomedical concepts, relationships and associated evidence. This work arises out of an earlier effort to develop a web application called “*Knowledge.Bio*” [7] to provide enhanced navigation through the knowledge base of PubMed cited concepts and relationships, captured by text mining in the Semantic Medline Database [8]. The knowledge harvesting workflow underlying *Knowledge.Bio* is here elaborated into a distributed web service network across diverse knowledge sources hosted within the NCATS Biomedical Data Translator Consortium, a publicly funded project supporting the FAIR integration of distributed biomedical research data and knowledge to accelerate the development of new disease treatments and reduce the barriers between basic research and clinical advances [9]. The outcome of this work was an iteratively refined web service specification implemented in an initial set of Beacons, with validation tools and client applications.

## Methods

The Knowledge Beacon API is a Swagger 2.0 specification that defines a set of endpoint paths embodying operations for accessing knowledge sources and discovering shared semantics for concepts and their relationships (Fig 1).

**Fig 1.**
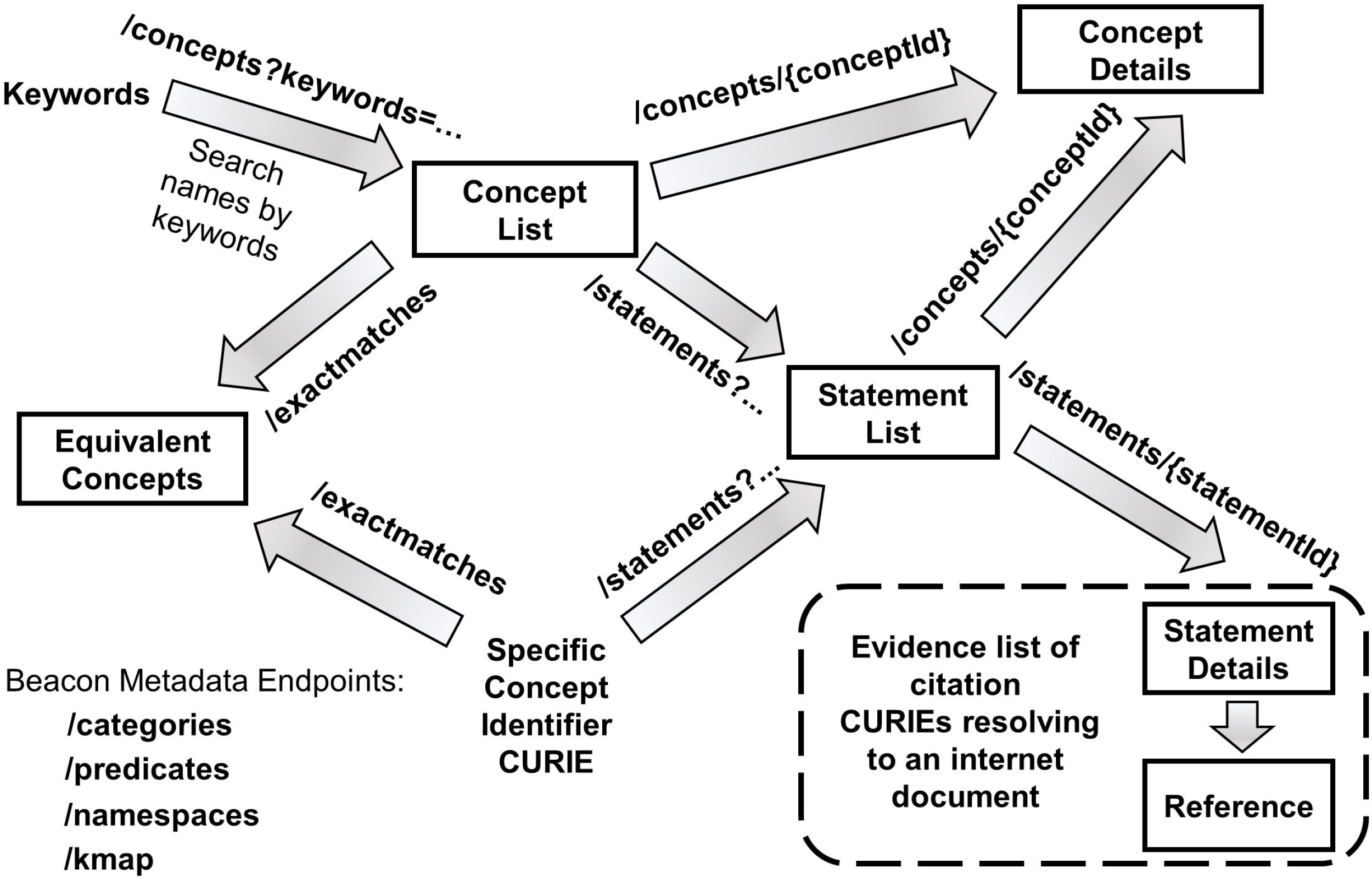
Knowledge Beacon Workflow. General step-by-step flowchart illustrating the sequential invocation of Beacon web service endpoints, with data flows as indicated. Also enumerated at the bottom left hand corner of the diagram is the set of metadata endpoints that report semantic terms and namespaces used by the Beacon in the annotation of results.

A Knowledge Beacon (hereafter abbreviated “Beacon”) initiates a workflow for knowledge discovery by simple search using a concept endpoint either with a *keywords* parameter (**/concepts?keywords=**) or one with a Compact Uniform Resource Identifier (CURIE; https://en.wikipedia.org/wiki/CURIE) of the concept (**/concepts/{conceptId}**). In both cases, one or more specific concepts with associated core details are retrieved.

Once identified, the canonical CURIE identifier of a chosen concept selected from the retrieved list is used as an input parameter to access a list of statements about the concept, documented as subject/predicate/object assertions (**/statements?s=…** where **s** is a **s**ubject canonical concept CURIE). Additional documentation, including supporting citations, associated with returned statements may be examined by calling the statement’s endpoint again with the statement identifier of one of the entries returned from the initial call (i.e. **/statements/{statementId}**).

The data model, concept data type (“*category*”) and relationship predicate (“*edge_label*”, “*relation*”) terms in results returned by a Beacon are compliant with an emerging public Biomedical Data Translator Consortium semantic standard and data model, the Biolink Model (**https://biolink.github.io/biolink-model/**). To assist client data parsing and interpretation, a Beacon supports several additional endpoints that return metadata summaries of Biolink Model terms specifically employed by the Beacon to annotate concepts and statements which are returned: concept type “categories” (**/categories**), identifier name spaces (**/namespaces**), relationship “predicates” (**/predicates**) plus a “knowledge map” of available subject-predicate-object triplet statement combinations (**/kmap**).

## Results

### Sample workflow

Knowledge Beacon workflows are implemented as a chained series of REST API endpoint calls that return data as JSON formatted documents, annotated using Biolink Model standards as noted above. Here we illustrate a basic minimal two step sequence of such calls which first identifies a list of concepts with names matching a keyword, then uses the identifier of one returned concept entry to retrieve *subject-predicate-object* statement assertions relating to that selected concept.

#### Step 1

Query knowledge sources by keyword to identify concepts. For example, calling the basic /concepts endpoint using the Fanconi Anemia complementation group C gene ‘FANCC’ as a keyword, on the Monarch “Biolink API” Beacon, namely:

~~~
https://kba.ncats.io/beacon/biolink/concepts?keywords=FANCC
~~~

returns the following JSON result with lists of CURIE-identified concepts (one entry shown; full list of concept entries truncated for conciseness):

~~~
                    [
                    … *some JSON results*
                    {
                       "categories": [
                         "gene",
                         "sequence feature"
                      ],
                      "id": "NCBIGene:102158362",
                      "name": "FANCC"
                     },
                     … *more JSON results*
                    ]
~~~

#### Step 2

Using the canonical (URL-encoded) concept CURIE of a selected concept in the list of concepts returned by keyword in step 1 above, e.g. NCBIGene:102158362, a search for knowledge assertions (statements) is made on the same database:

~~~
https://kba.ncats.io/beacon/biolink/statements?s=NCBIGene%3A102158362
~~~

This query gives another JSON result which contains asserted “subject-predicate-object” statements about the concept, where the predicate return defines the relationship, as follows:

~~~
                    [
                      {
                        "id": "biolink:125a0182-0205-44a8-a70a-c03339383177",
                        "object": {
                          "categories": [
                            "gene"
                          ],
                          "id": "NCBIGene:102158362",
                          "name": "FANCC"
                      },
                      "predicate": {
                          "edge_label": "is_about",
                          "relation": "IAO:0000136"
                      },
                      "subject": {
                          "categories": [
                            "publication"
                          ],
                          "id": "PMID:17145712",
                          "name": "PMID:17145712"
                      }
                     },
                     … more JSON results
                    ]
~~~

### Beacon implementations

A stable set of publicly accessible Beacons are implemented and currently hosted stably online (as of February 2020) by the NCATS Biomedical Translator Consortium, as enumerated in Table 1. The Java and Python software implementations of these Beacons are available in repositories of the NCATS-Tangerine (https://github.com/NCATS-Tangerine) GitHub organization. One implementation is a generic accessor of Biolink Model compliant knowledge graph databases stored in Neo4j (https://github.com/NCATS-Tangerine/tkg-beacon). These Beacon implementations may be tested using an available validator application (https://github.com/NCATS-Tangerine/beacon-validator). A Python command line Beacon client is available (https://github.com/NCATS-Tangerine/tkbeacon-python-client). A Knowledge Beacon Aggregator (https://github.com/NCATS-Tangerine/beacon-aggregator-client) was also designed to manage a registered pool of Beacons, and to return consolidated knowledge using “equivalent concept cliques” to merge related Beacon results.

**Table 1.**
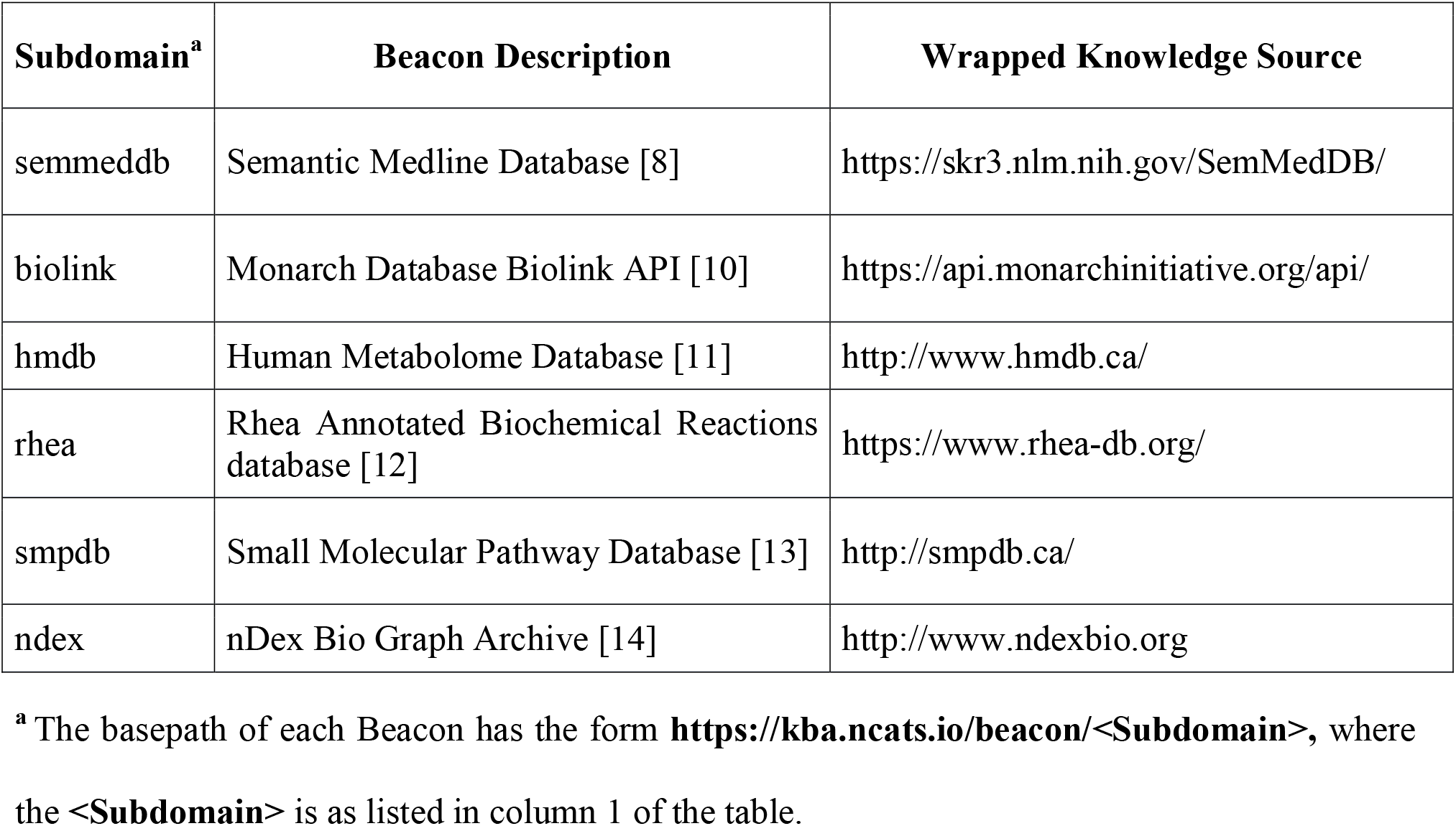
Biomedical Translator Consortium Deployed Beacons.

| Subdomain<sup>a</sup> | Beacon Description | Wrapped Knowledge Source |
| --- | --- | --- |
| semmeddb | Semantic Medline Database [8] | https://skr3.nlm.nih.gov/SemMedDB/ |
| biolink | Monarch Database Biolink API [10] | https://api.monarchinitiative.org/api/ |
| hmdb | Human Metabolome Database [11] | http://www.hmdb.ca/ |
| rhea | Rhea Annotated Biochemical Reactions database [12] | https://www.rhea-db.org/ |
| smpdb | Small Molecular Pathway Database [13] | http://smpdb.ca/ |
| ndex | nDex Bio Graph Archive [14] | http://www.ndexbio.org |

<sup>a</sup> The basepath of each Beacon has the form **<https://kba.ncats.io/beacon/<Subdomain>>**, where the **<Subdomain>** is as listed in column 1 of the table.

## Discussion

The Knowledge Beacon API is a basic knowledge discovery workflow (Figure 1) representing a relatively high-level use case of user interaction with the biomedical knowledge space, and as such, lacks the full expressive power of a general knowledge query language interface like SPARQL. Furthermore, the beacon data model aligns with the emerging Biolink Model standards of the Biomedical Translator Consortium as its template for knowledge representation. As such, Beacons do not automatically express results in a generic manner as do knowledge representations such as RDF, although conversion of Beacon statement results into RDF format is easily accomplished. Finally, aside from some general profiling of the performance of the Swagger API endpoints on various knowledge sources, we have not here conducted a rigorous computing-theoretic assessment of the efficiency of this form of knowledge harvesting, although early experience with Beacons point to challenges with internet latency and knowledge-source specific differences in query performance. In partial response to such challenges, we prototyped a “Knowledge Beacon Aggregator” to provide enhanced asynchronous query/status/retrieval endpoints as a client-friendly integration layer for managing access to, and merging data from, a registered catalog of multiple Beacon implementations.

Despite the use of some off-the-shelf API generation tools, the wrapping of knowledge sources as Beacons remains a labour-intensive activity. The semantics of the knowledge source being wrapped must be heuristically translated. This is somewhat easier for knowledge sources which have a small number of easily resolved discrete data types (i.e. discrete Biolink Model concept categories of data) and namespaces with clear mapping onto those discrete data types.

In contrast, some “graph” knowledge sources, for example, the NDex Bio biomedical network data archive (https://home.ndexbio.org/index/, wrapped by this project as the *ndex* Beacon), don’t have such clear concept category and relationship predicate tagging of much of the archived data. The development of useful but (so far) imprecise heuristics to tag such data on the fly is required to develop a useful Beacon. In other cases, such as biomedical knowledge resources whose data object namespace aggregates several types of concepts in a fuzzy manner with limited additional concept category tagging, it may be even more challenging to semantically tag data entries for beacon export.

A few common library and reference implementations are developed for Beacons; the Beacon platform would benefit from the further development of standardized tools to systematically assist such wrapping of native knowledge sources.

The availability of a shared API standard for knowledge integration doesn’t, in and of itself, deal with all the challenges of FAIR data integration within the global community. Practical experience with knowledge harvesting using such API implementations has revealed performance issues relating to internet and service latency, bandwidth limitations. Knowledge warehousing in centralized knowledge graphs using ETL (Extract, Transform, Load) processes may sometimes result in a more tractable process for biomedical knowledge integration; however, such approaches are still faced with the task of merging equivalent concepts, including the elimination of duplicate concepts and the resolution of conflicting information, including weighting of assertions differing in levels of confidence. More unique to ETL warehousing approaches is the ongoing problem of keeping such resources up-to-date relative to their original knowledge sources. Note that ETL warehouses and API driven distributed knowledge harvesting approaches can be complementary, in that ETL data warehouses can also themselves be accessed by the application of web service REST API’s like the Knowledge Beacon API. In fact, some of the current Beacon implementations use this approach: a back end Biolink Model compliant Neo4j knowledge graph directly wrapped with the API.

The Linked Open Data paradigm using RDF knowledge representation and SPARQL represents an alternate paradigm for distributed knowledge integration, the theoretical performance of which was surveyed by Verborgh *et al* [6]. In their assessment, it was noted that downloadable RDF knowledge data sets and SPARQL endpoints to triple store knowledge bases represent two extremes of a continuum of RDF knowledge access, each with their characteristic advantages and weaknesses. They proposed that a constrained query selector specification and RDF representation – with data, metadata and hypermedia controls - denoted as Linked Data Fragments could be a shared design representation spanning both ends of the continuum. Furthermore, they proposed an intermediate implementation - termed Triple Pattern Fragments - partitioning RDF processing more symmetrically across client and server, thus potentially mitigate some of the challenges of both ends of the API design continuum, for more balanced client-server performance and greater ease of implementation (see https://github.com/LinkedDataFragments).

Generally, API approaches to knowledge harvesting may work best with use cases involving smaller batches of knowledge retrieval based on a focused navigation of the knowledge space from larger open-ended data sources which would be refractory to import into centralized knowledge graphs.

Finally, there are two other API standards of the Biomedical Data Translator Consortium: the “NCATS Reasoner API” (https://github.com/NCATS-Tangerine/NCATS-ReasonerStdAPI) and the Biothings API (https://biothings.io/). Although there are parallels between them, the Beacon API is a simpler lower level interface to knowledge resources than the Reasoner API, and is somewhat more constrained to the Biolink Model than the Biothings API. But the utility of the Reasoner API is inspiring efforts to publish a Reasoner API interface on top of an implementation of Knowledge Beacons (https://github.com/NCATS-Tangerine/kba-reasoner). We did also implement prototype Beacon wrapper for the Biothings API (https://github.com/NCATS-Tangerine/biothings-explorer-beacon).

## Availability

Knowledge Beacon software is open source licensed and available for access in GitHub. A suitable introduction to the API, containing references to related software components, can be found at **https://github.com/NCATS-Tangerine/translator-knowledge-beacon**.

## Acknowledgments

BMG and RMB collaborated on the predecessor “Knowledge.Bio” application embodying the workflow captured by the Knowledge Beacon API. CJM and BMG coined the name “Knowledge Beacon” to express the architectural vision of uniformly wrapped knowledge sources for distributed knowledge discovery and harvesting. RMB and his team elaborated the original software design of web service endpoints and initial code implementations embodying Beacon functionality, then guided further iterations of the API based on feedback from colleagues within the Biomedical Translator Consortium, with special mention to co-author VD who proposed insightful revisions to the API, based on his direct experience implementing a Beacon to wrap HMDB. Under overall supervision by RMB, the heavy lifting of iterative software development of Beacon implementations - including several beacons, client (including aggregator) and validation applications - was undertaken by LMH while he was a member of the STAR Informatics team, assisted by the valuable software programming contributions of several computing science cooperative education students: MG, WTS and KCHB.

The authors would like to sincerely thank Nomi Harris and Marcin Joachimiak of LBNL for their very helpful editorial feedback on, and suggested revisions to the draft manuscript.

The authors would also like to thank the various members of the Biomedical Data Translator Consortium who gave helpful user needs feedback and support of the Knowledge Beacon API during its development, in particular, Chris Bizon and Stephen Ramsay. We also acknowledge here Greg Stupp who, while employed at TSRI, implemented an earlier version of a Beacon wrapper for biomedical knowledge in Wikidata.

